# An evaluation model for aboveground biomass based on Hyperspectral Data from field and TM8 in Khorchin grassland, China

**DOI:** 10.1101/792424

**Authors:** Xiaohua Zhang, Meirong Tian, Xiuli Chen, Yongjun Fan, Jianjun Ma, Danlu Xing

**Author notes:** These authors are co-first authors on this work (Xiaohua Zhang, Meirong Tian, Xiuli Chen).

## Abstract

Biomass is an important indicator for monitoring vegetation degradation and productivity. This study tests the applicability of Hyperspectral Remote-Sensing in situ measurements for high-precision estimation aboveground biomass (AGB) on regional scales of Khorchin grassland landscape in Inner Mongolia, China. Field experiments were carried out which collected hyperspectral data with a portable visible/NIR hyperspectral spectrometer (SOC 710), and collected aboveground net primary productivity (ANPP). Ground spectral models were then developed to estimate ANPP from the normalized difference vegetation index (NDVI), which was measured in the field following the same method as that of the Thematic Mapper(TM) from the Landsat 8 land imager (TM_NDVI). Regression analysis was used to assess the relationship between ANPP and NDVI based on coefficients of determination (R^2^) and error analysis. The estimation of ANPP had unique optimal regression models. By comparing the different spectral inversion models, we selected an exponential model associating ANPP with NDVI (ANPP = 12.523*e3.370*(0.462*TM_NDVI+0.413), standard error = 24.74 g m-2, R^2 =^ 0.636, P < 0.001). This study suggests that the model can be used to monitor the condition and estimate the productivity of grassland at regional scales. The results still show a high potential to map grassland degradation proxies on the ground hyperspectral model. Thus, this study presents biomass hyperspectral inversion technology to remotely detect and monitor grassland degradation and productivity at high precision.

## Introduction

The tools for remotely sensing of vegetation have evolved significantly in recent decades[1], and spectral imaging has become increasingly popular in remote-sensing research for correlating spectral data with the biophysical properties of vegetation. Hyperspectral remote-sensing data have subsequently been widely used to estimate vegetation biomass[2–5], vegetation cover (VC)[6–7], nitrogen content[8], and the leaf area index of vegetation[1,9].

The accurate estimation of aboveground net primary productivity (ANPP) is an active area of research and can provide valuable information about the productivity and ecosystem service value of grassland [10]. ANPP is an important impact factor for desertification and are often used as indicators for monitoring and evaluating grassland productivity and degradation [11]. The present study aimed to develop models for estimating biomass and VC based on satellite data, which allow an assessment over large areas at a low cost [12]. Desertification in the Khorchin grassland is becoming worse with the rapid expansion of the population, overgrazing and the warmer and drier trend associated with climate change [13]. The accurate estimation of the biomass of the grassland over large areas using remotely sensed data is thus very important for monitoring desertification and for improving the scientific management of grassland ecological resources.

Remote-sensing data have been transformed and combined into various spectral vegetation indices that are used as predictors of parameters, such as the normalised difference vegetation index (NDVI), ratio vegetation index, perpendicular vegetation index, soil adjusted vegetation index, and transformed soil adjusted vegetation index [14–17]. Most studies apply NDVI because it minimises the effects of topography[18] and is more reliable for the estimation of biomass of ecosystems/habitats dominated by grasses when the grasses are actively growing [9,19]. The NDVI is thus widely used to characterise grass ecosystems and to estimate biomass and VC[3,4,19–23].

NDVI have been determined from data sets collected by various satellite instruments, such as the Landsat Thematic Mapper (TM), the National Oceanic and Atmospheric Administration/Advanced Very High Resolution Radiometer (NOAA/AVHRR), MODIS, Gaofen-2 and the Moderate Resolution Imaging Spectro radiometer[9,24]. The Landsat 8 with narrowband indices are highly suitable to be chosen to map AGB accurately, because the narrowband indices have led to significant improvements in the predictive capability of models, and hyperspectral data from aerial imagery or field spectrometry have the potential to estimate the biophysical properties of rangeland or steppe vegetation with a greater accuracy than broadband indices [3,4,23,25–26].

In order to improve the accuracy of biomass estimation, the measurements of ground reflectance have been used to estimate biomass in grasslands and steppes since the 1970s [27], but ground spectral reflectance can be influenced by variable factors of the landscape such as the distribution of plant communities [28], soil colour [29], hydrology[30], and topography[31], and sensor radiance may be strongly affected by atmospheric scattering[32]. For these reasons, various regression models have been established for associating vegetation indices with biomass in different areas. Relationships have been established between remote-sensing data and the biophysical properties of vegetation, mostly linear or non-linear, that have greatly improved the accuracy of biomass estimates and that have determined patterns of grassland productivity in various regions [6,33–37].

Taking into account the advantages and disadvantages of the current remote sensing sources to estimate AGB, In this study, we present a remote sensing approach for estimating and monitoring AGB in meadows and pastures during the growing season. We used remote sensing of Landsat 8 and ground hyperspectral to calculate the normalized difference vegetation index (NDVI), and on-field aboveground net primary production (ANPP) measurements to establish an empirica1 exponential model to estimate spatial ANPP across the entire Khorchin grassland.

The main objectives of this study are: (i) to analyse the relationship of the ground narrowband NDVI with ANPP and then to develop the most suitable ground spectral models for evaluating ANPP over a large area of the Khorchin steppe, and (ii) to use the ANPP model to understand the biocapacity of the Khorchin grassland for providing technical support for determining a reasonable grazing intensity and guiding the development of the livestock industry.

## Materials and methods

### Study area

The Khorchin grassland is located in the eastern section of the ecotone between crop production and animal husbandry, we choose the region of Bairin Youqi in Inner Mongolia of China as the study area (Fig. 1), which is an important component of the Khorchin grassland and is typically sensitive and fragile. Bairin Youqi has a semiarid, temperate, continental monsoon climate with mean annual temperature of 4.9 °C and mean annual precipitation of 358 mm (precipitation is less than evaporation). From low to high elevations, the distribution of vegetation is meadow, sandy vegetation, and low mountain grassland respectively. The dominant grassland species include *Achnatherum splendens* (*Trin.*) Nevskia, *Stipa capillata* Linn., *Leymus chinensis (Trin.)* Tzvel., and *Agropyron cristatum (Linn.)* Gaertn.

**Fig. 1.**
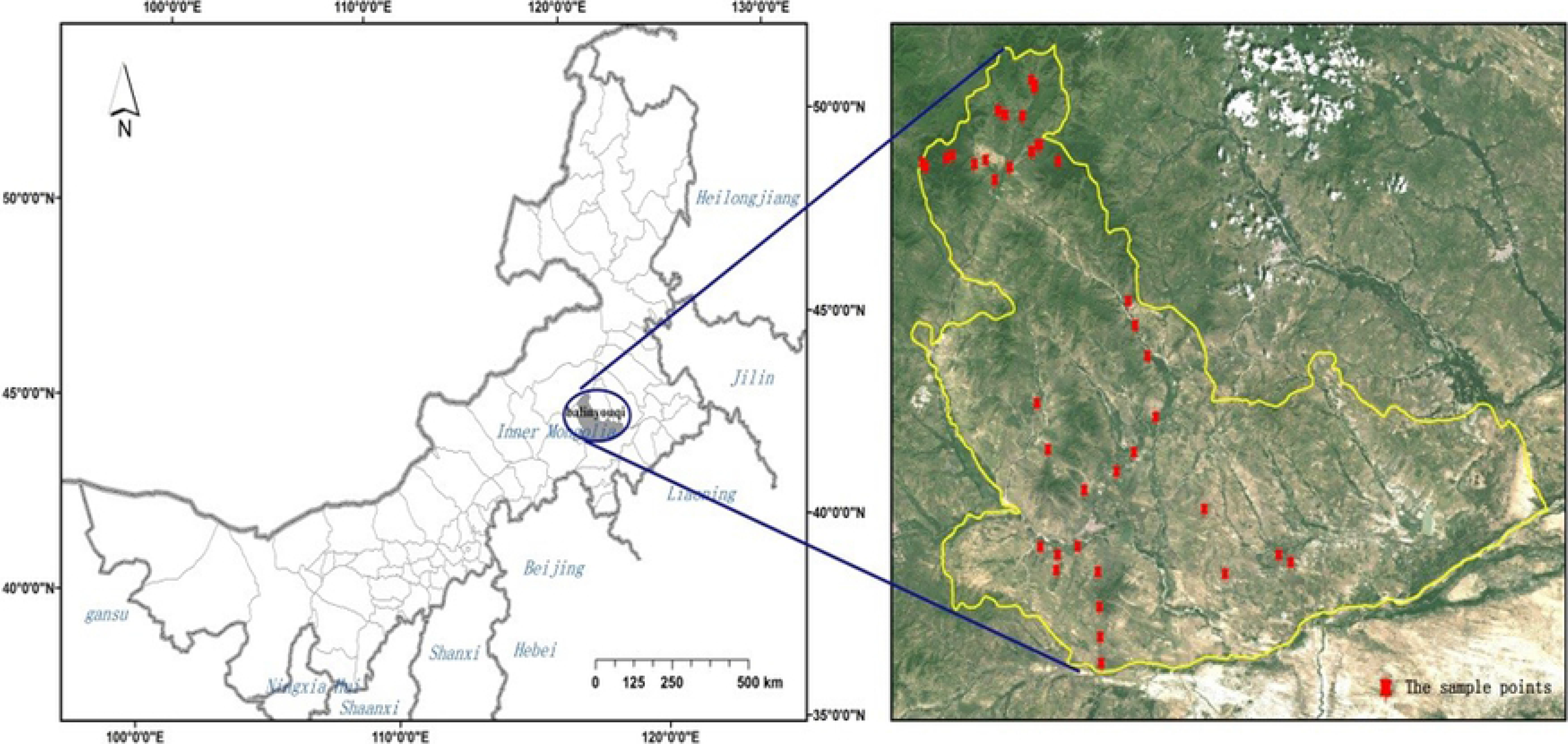
Map of Inner Mongolia (left) and the location of the sampling sites in Bairin Youqi (right) This study was carried out in the field of Khorchin grassland which was State-owned Land and did not involve endangered or protected species. Meanwhile, because this study supported by National Environmental Conservation Research Program, so the government of Bairin Youqi permitted and approved this study.

## Collection of field data

### Experimental setup

Field work was conducted during 15-30 July 2016, coinciding with the most productive period of vegetational growth. Based on the topography and land use, 39 plots were established that included large, homogeneous patches of vegetation and representative vegetational communities with different types of vegetation. The plot size was set at 30 × 30 m, equivalent to the size of a TM8 pixel. The plots contained a total of 173 quadrats of 1 × 1 m. The data collected were divided into 2 groups. Group one, which contained 153 quadrats were used to build the ground spectral model; group two, which contained 20 quadrats, were used for the accuracy test of the spectral inversion model. Meanwhile, within group one, the data of approximately two thirds of the total quadrats (n=115) were chosen randomly to build the model while the rest were used for testing the terrain model in terms of selecting the best fitting function and precision.

### Field spectral data

The field data were collected using the SOC710 Hyperspectral Imaging System which Manufactured by Surface Optics Corporation in America. The SOC710 is a precision instrument with an integrated scanning system and analysis software that can quickly obtain high-quality hyperspectral images at visible to near-infrared (NIR) wavelengths in the range 0.4-1.0 μm. The system can be used under normal lighting conditions at variable exposures and gains. The SOC spectra were collected with a 10° field of view and at 1.2 m above the grass canopy. All spectral measurements were collected between 9:00 and 15:00 Beijing time under clear skies. Three measurements were taken for each sample of grass canopy. These spectra were standardised to spectra measured at approximately 10-minute intervals with a white board. The average of three replicates for each sample was used for the analysis.

### Biomass measurements

After the spectral data had been recorded, the standing biomass was collected in the quadrats at each sample location. The fresh weight of green herbaceous material was recorded soon after clipping, the samples were then dried at 80 °C for 10-12 hours, and the dry masses of the samples were determined.

## Image data acquisition, satellite data, and preprocessing

Biomass was assessed using TM8 data from the Landsat 8 land imager of the United States Geological Survey. The satellite data were acquired within the same time frame in which the field data had been collected, and the images were free of clouds and haze. Four suitable TM8 satellite scenes at PATH/ROWs 123/29, 123/30, 122/29, and 122/30 were analysed. The satellite data were geometrically rectified by a digital elevation model and ground-control points from Land Survey. The four TM8 scenes were processed for atmospheric correction with the Fast Line-of-sight Atmospheric Analysis of Spectral Hypercubes software package.

## Data analysis

### NDVI calculation

NDVI are commonly calculated from RED and NIR reflectance data [38]. We calculated the SOC_NDVI of the samples from SOC710 spectral reflectance using the ENVI 5.0 image analysis software. The method for calculating NDVI was the same as that used for calculating the TM_NDVI:

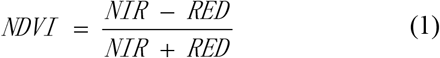

Where the RED and NIR bands correspond to wavelengths of 630-680 and 845-885 nm, respectively. Spectral reflectance data should be resampled within the scope of the RED and NIR bands.

### Regression analyses

The regression analyses were carried out for the scatter diagrams of ANPP vs. SOC_NDVI, and SOC_NDVI vs. TM_NDVI. In study area, The 173 quadrats data were employed to obtain the regression model for ANPP vs. SOC_NDVI. Mean value of NDVI within a specific plot was calculated, and then the data of the total 39 plots (Fig. 1) were used in the regression analysis for SOC_NDVI vs. TM_NDVI.

The coefficient of determination (*R*^2^) and the adjusted *R*^2^ were used to test the strength and significance of the relationships between the field data and the corresponding data extracted from the satellite scenes. The standard error (SE, Eq. 2) of the prediction based on the independent test data and the coefficient of mean error (MEC, Eq. 3) were calculated to assess the accuracy of the developed models.

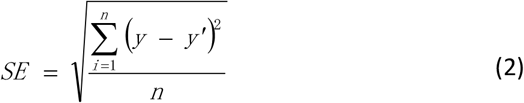

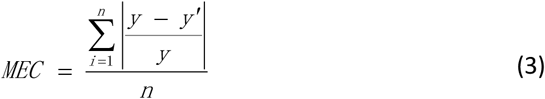

where y is a measured biomass, y’ is an estimated biomass for the test data, and n is the number of samples.

## Results

### Optimal ground spectral models and tests of model accuracy

#### The optimal ground spectral models for biomass

From the analysis and evaluation of the relationships between ANPP and SOC_NDVI computed from reflectance data obtained by the SOC710 in the field, we chose linear, logarithmic, power, and exponential functions to fit and optimise the regression equations for selecting the best regression model (Fig.2).

**Fig.2.**
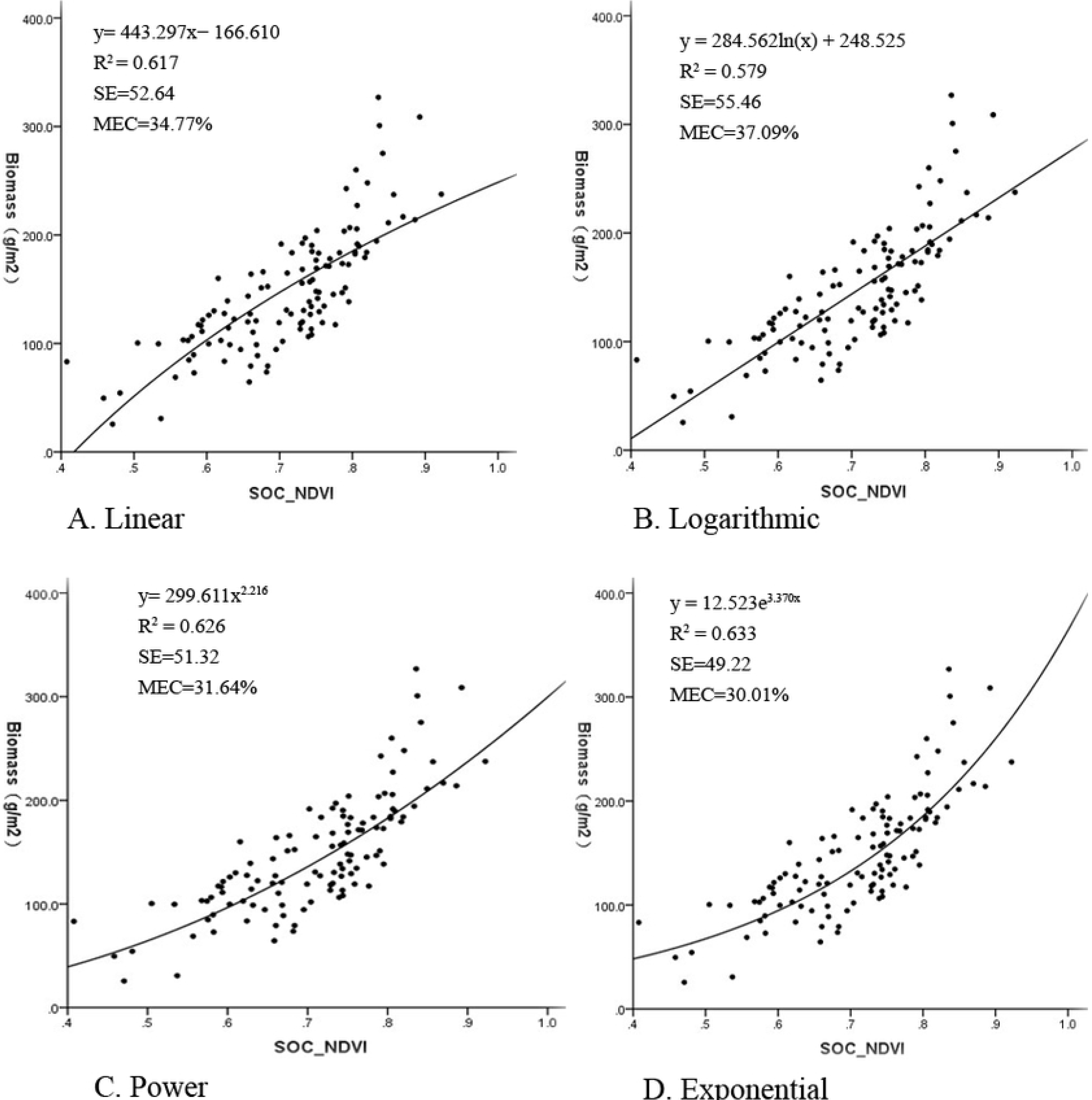
The Simulation Curves of the Regression Equation of the Training Samples. The relationships between ANPP and SOC_NDVI was significant (*P* < 0.001) for all functions and met the assumptions of the statistical analyses. The exponential model was superior for ANPP, with an *R*^2^ of 0.636, indicated by bold type in Tables 1.

**Table 1.** Comparison of the regression equations between ANPP and SOC_NDVI.

|  | Linear | Logarithmic | Power | Exponential |
| --- | --- | --- | --- | --- |
| $n$ | 115 | 115 | 115 | 115 |
| Equation | $y = 443.297x - 166.610$ | $y = 284.562\ln(x) + 248.525$ | $y = 299.611x^{2.216}$ | $y = \mathbf{12.523e^{3.370x}}$ |
| $R^2$ | 0.617 | 0.579 | 0.626 | **0.636** |
| Adjusted$R^2$ | 0.614 | 0.575 | 0.623 | 0.633 |
| $F(\alpha=0.01)$ | 182.255 ($P < 0.001$) | 155.181 ($P < 0.001$) | 802.5746 ($P < 0.001$) | 1089.7635($P < 0.001$) |

Note: $n$ , number of samples; $R^2$ , coefficients of determination; the best regression model is highlighted in bold.

#### Tests of model accuracy

The accuracy of the models was tested to obtain the best regression models. We used test sets of all field samples to analyse and evaluate the errors in the regression models (Table 2). A comparison of the predictive performances of the regression equations indicated by SE and MEC are presented in Table 2.

**Table 2.**
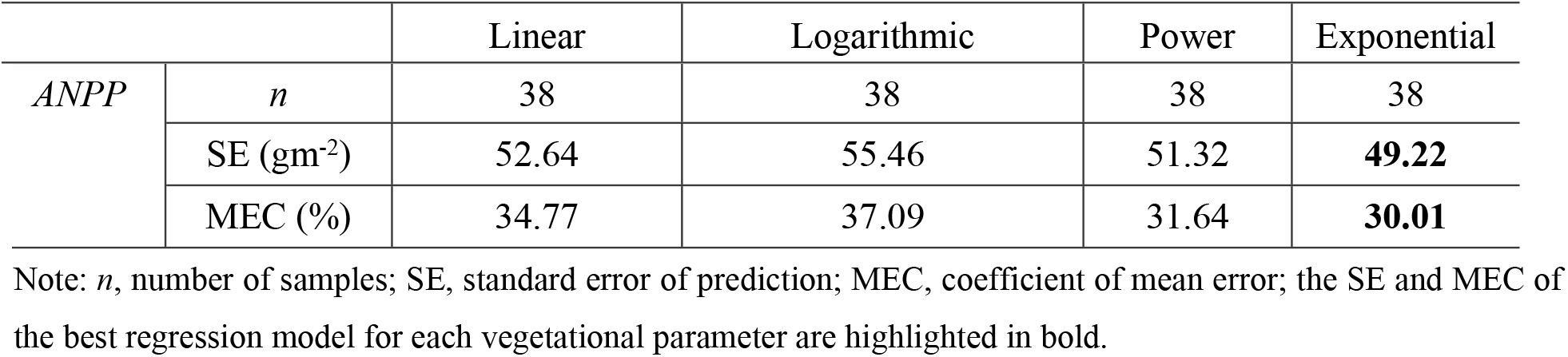
Comparison of the errors of the regression equations.

**Table 3.** Regression equations between TM_NDVI and SOC_NDVI.

| Equation | *n* | *R* | $R^2$ | $F (\alpha=0.01)$ |
| --- | --- | --- | --- | --- |
| $y=0.462x+0.413$ | 39 | 0.810 | 0.656 | 70.63, $P < 0.001$ |

Note: *n*, number of samples; *R*, $R^2$ , coefficients of determination.

We determined the best models for ANPP based on *R*^2^ and the independent validations. The exponential equation was optimal for ANPP (*R*^2 =^ 0.636, SE = 49.22gm^−2^, MEC = 30.01%; Tables 1 and 2). Models with the following equations (Eqs. 4) were selected and used as the optimal ground spectral models for ANPP of the entire Khorchin grassland:

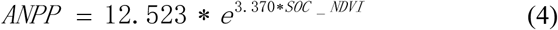

### The relationship between TM_NDVI and SOC_NDVI

The linear regression equation was selected based on the analysis of the TM_NDVI/SOC_NDVI scatter plot. The relationship between TM_NDVI and SOC_NDVI was significant, with an *R*^2^ of 0.656 (*P* < 0.001) and met the assumptions of the statistical analyses (Table 4, Fig.3). The model with the following equation (Eq. 5) was selected for the relationship between TM_NDVI and SOC_NDVI for the entire Khorchin grassland:

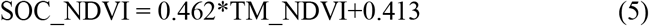

**Fig. 3.**
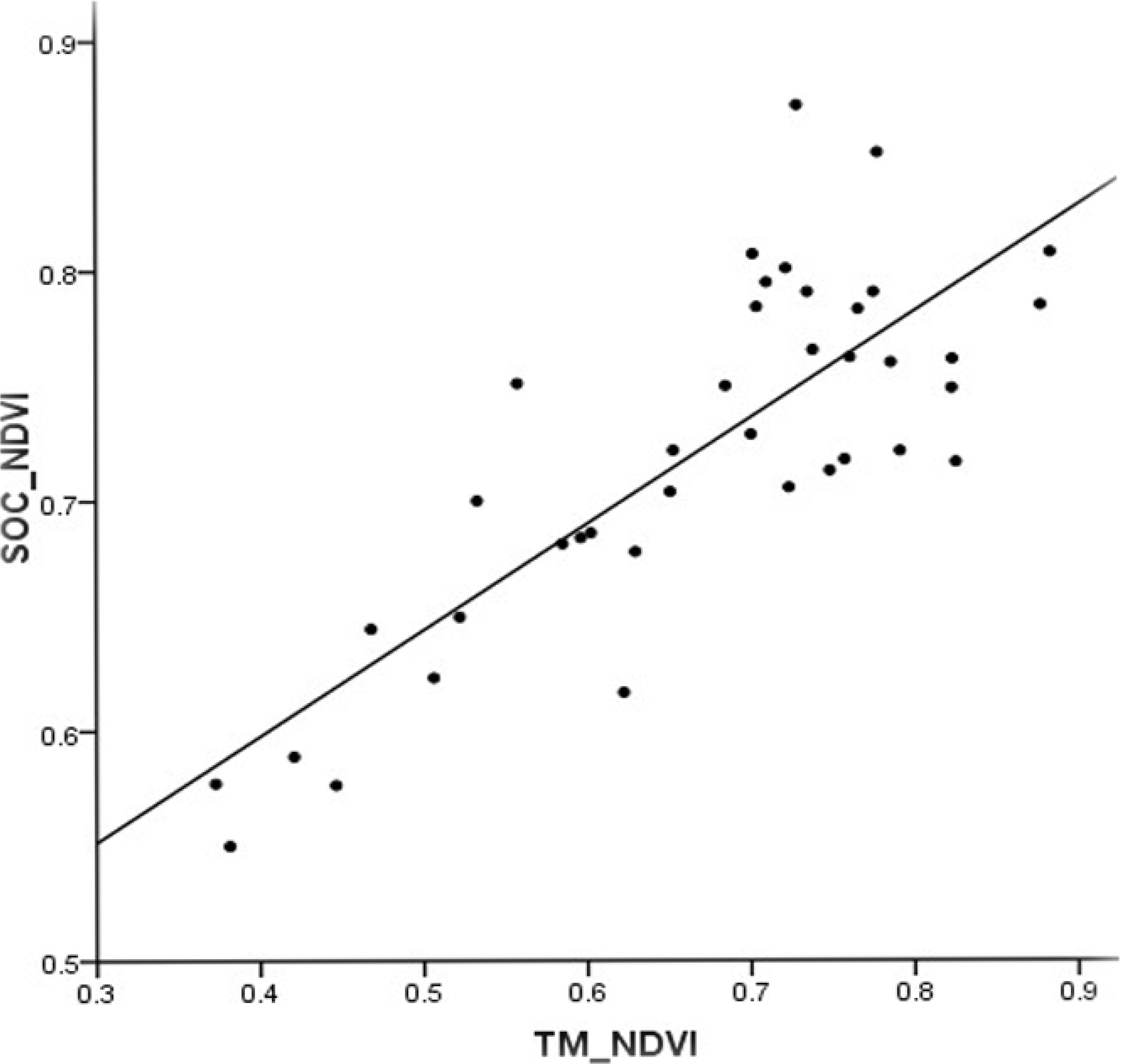
Fitted curve of the best model for the relationship between SOC_NDVI and TM_NDVI.

### Spectral inversion models

The spectral inversion models of TM8 for ANPP was calculated by Eqs. 4–5:

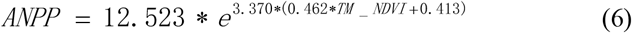

To test the agreement between measured and predicted values, we applied Eqs. 6 to the TM8 NDVI greyscale image and obtained the patterns of ANPP distribution in the study area by grid computing. The test data sets were then converted into vector diagrams defined by geographic coordinates by geographic information system. The values at the test points were recorded in the distribution patterns as the corresponding pixels predicting values of ANPP. The relationship between actual and predicted values was used to evaluate the accuracy of model.

The correlation between the predicted and actual values was significant, as were the independent validations for predicting biomass (SE = 24.74, MEC = 18.61%; Fig.4). This study suggested that the spectral inversion models could be used to monitor grassland biomass at regional scales.

**Fig. 4.**
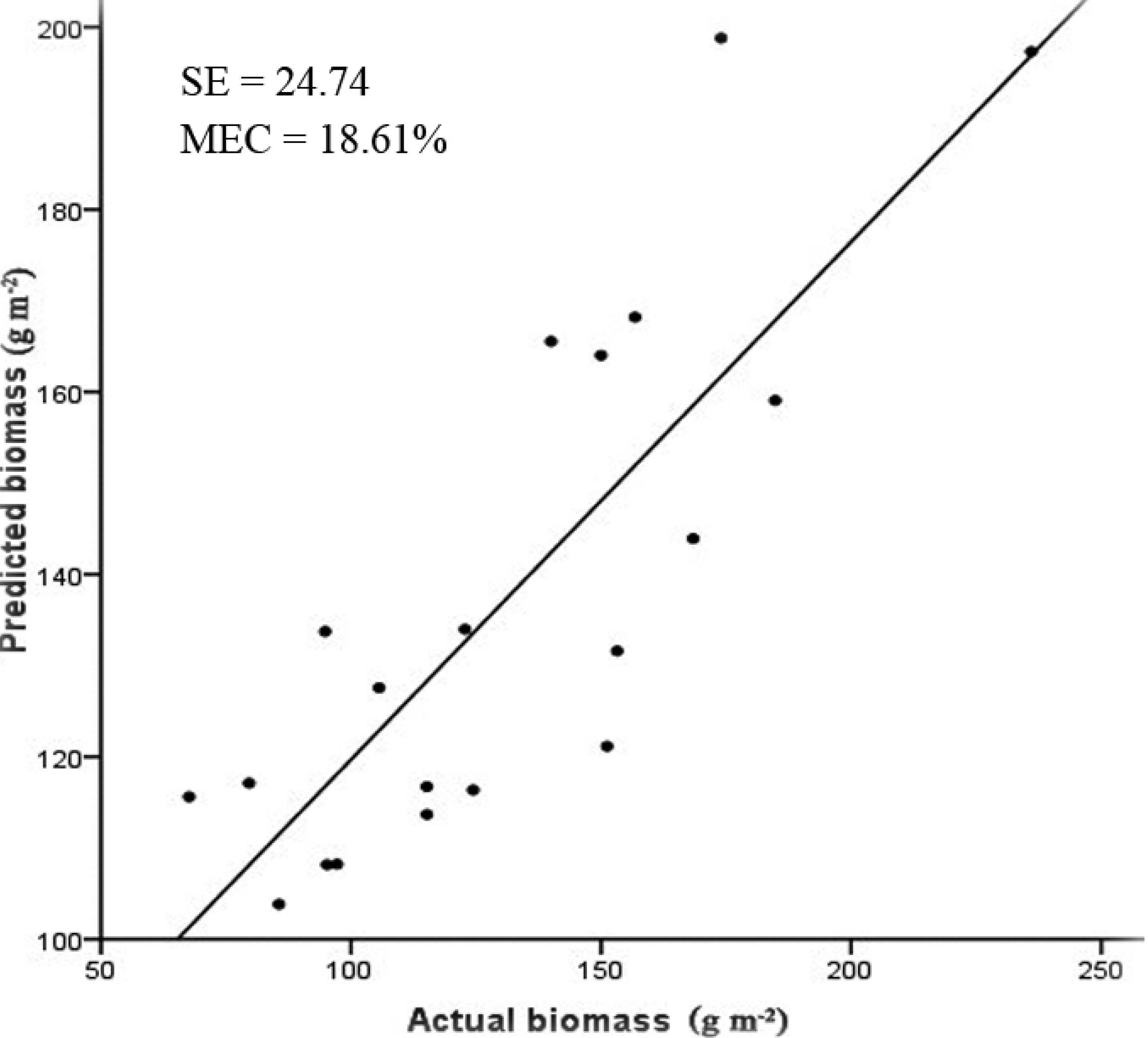
Independent validation for predicting biomass (*n* = 20 *P* < 0.001). SE, standard error of predicted biomass; MEC, coefficient of mean error.

## Discussion

The main goal of this study was to establish more accurate models for estimating ANPP of the grassland in Khorchin. We chose the NDVI vegetation index, which can be calculated from spectral reflectance data acquired in the field and from data from Landsat TM7 Band 4 (TM4; 760-900 nm) and Band 3 (TM3; 630-690 nm) or from NOAA/AVHRR Channel 1 (580-680 nm) and Channel 2 (720-1100 nm)[39–41]. In a previous study, we also calculated the NDVI from data collected by a FieldSpec3 spectroradiometer (Analytical Spectral Devices, Boulder, USA), at spectral reflectances of 620-670 (RED) and 841-876 (NIR) nm [9]. To further improve the accuracy in the present study, we chose satellite data from Landsat 8(TM8), which have a higher geometric precision and signal-to-noise ratio than the other Landsat data, and used the SOC710 Hyperspectral Imaging System, which is more accurate than the FieldSpec3 spectroradiometer. The TM8 remotely sensed imaging data were only released in 2013, so they have not yet been widely applied to monitor vegetational biomass. This study applied the field data for monitoring the vegetation, thereby providing an informational baseline for this study area. The spectral inversion model was ideal, indicating that TM8 remote imaging can be used for research on vegetation biomass on a regional scale.

ANPP have their own optimal regression models based on the processing and statistical analysis of experimental data in the study area. The optimal equations for the estimation of ANPP (Fig.5) indicate that the relationship between SOC_NDVI and ANPP weakens at biomass >350 g m^−2^ for grassland. Estimates of biomass above these levels are inaccurate or unreliable and may be affected by the NDVI lower saturation phenomenon in areas of dense vegetation cover. When biomass exceed these levels, factors such as grass height and leaf area index must be considered, or a modified SOC_NDVI should be derived[42].

**Fig. 5.**
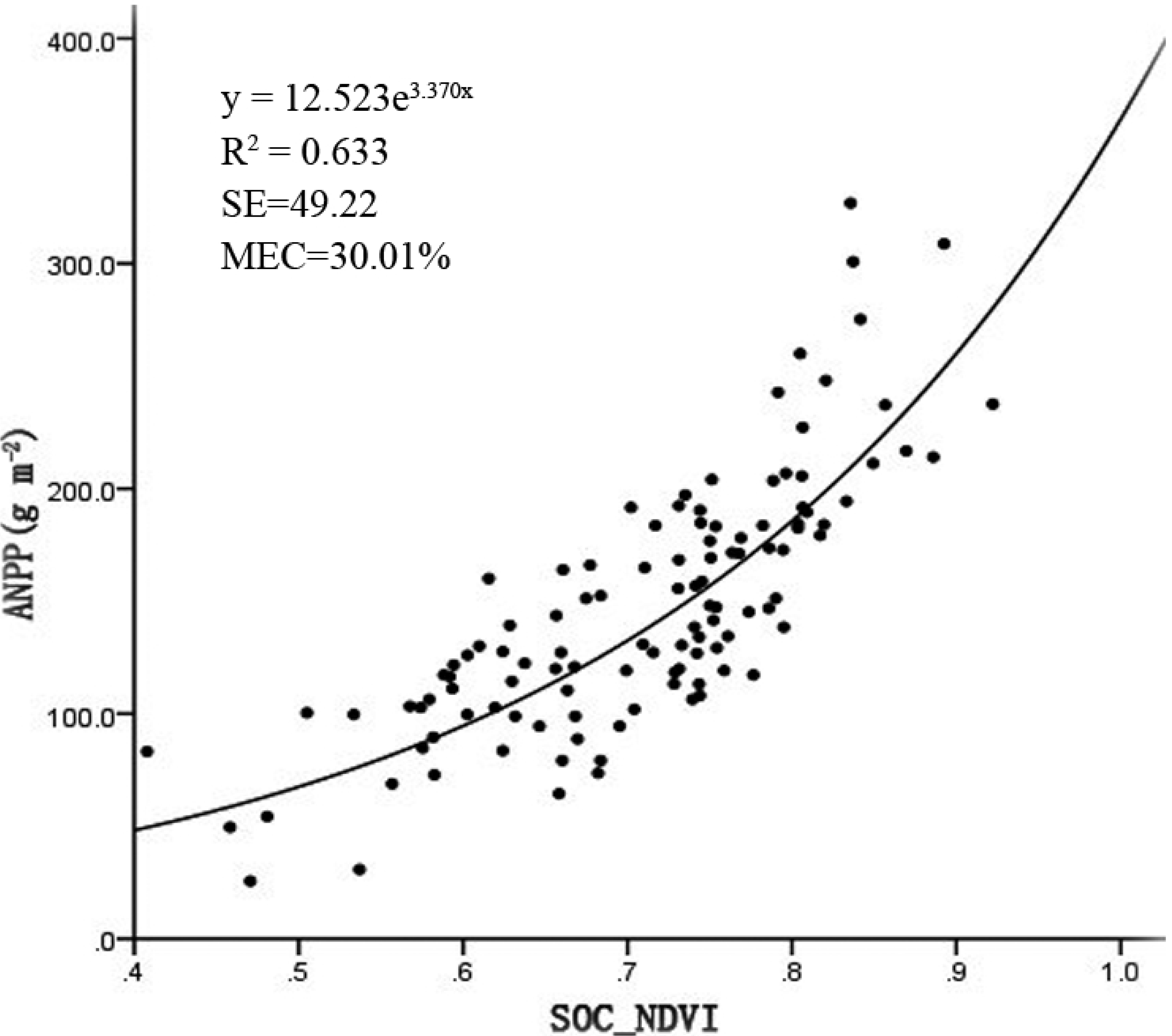
Fitted curve of the best model for the relationship between ANPP and SOC_NDVI for the calibration sets.

The ground spectral models for ANPP can be applied to TM8 images, because measured spectral characteristics of plants on the ground are intrinsically linked to those obtained by TM8 remote sensing. Grassland yield over large areas can be estimated based on the ground spectral model. The models, however, could be more accurate if field and satellite data are collected over several years rather than only for one year. Also, the field and satellite data should be acquired at the same time for maximal correspondence. In future field experiments, we will assess the collective influence of these vegetational characteristics and the NDVI on biomass prediction and will seek to obtain a modified NDVI for estimating the biomass of dense vegetation under natural conditions.

## Conclusions

This study developed a relatively accurate model for estimating AGB and tests the applicability of hyperspectral data from field and TM8 to map AGB on regional scales by a regression analysis method. The methodology we adopted in the study was a first attempt to Retrieval of vegetation biomass from ground hyperspectral remote sensing in Khorchin grassland.

The accuracy of ground spectral inversion is affected by many factors, and the quality of the selected remote sensing image data has the greatest impact on the fitting accuracy of the model. Landsat 8 satellite data is selected for remote sensing data, which has higher geometric accuracy and signal-to-noise ratio than previous Landsat data, which effectively expands the application range of image data. In the aspect of imaging mode, the sweep pendulum design of OLI imager has good stability and improves the image quality, and in the aspect of geometric accuracy, L1T data product is a data product after precise correction, and the product accuracy has been greatly improved. In this paper, TM8 data is used to retrieve vegetation biomass, and the results show that calculated *R*^2^ and SE and MEC values for various regression models vary among ground spectral models. By comparison, the exponential regression models we developed show a stronger relationship between spectral reflectance and ANPP. An exponential equation was optimal for estimating ANPP in the Khorchin grassland. Accuracy verification indicated that the relationship between the actual and predicted biomass was significant. Estimating ANPP with high accuracy based on NDVI derived from TM8 satellite data is thus possible, which accumulates experience for the application of TM8 data in vegetation monitoring field.

The accuracy of this technique depends on living, green biomass and not on senesced or dead biomass, so the timing of the acquisition of NDVI data is critical, and the model can possibly be improved if models are developed per vegetation types and using a larger range of ground data. In brief, this research shows the usefulness of hyperspectral data from field and TM8 to evaluate aboveground biomass at very high precision to provide theoretical and data support for RS monitoring, grassland governance and ecological restoration.

## Acknowledgements

We would like to thank Xueping Pan, Kun Wang, and Maorong Tian for their help with the collection and identification of the grasses, and thank Jixi Gao and Yanmei Chen for their help with guiding techniques and methods.

